# Engineering *Pseudomonas putida* KT2440 for open-loop upcycling of mixed plastics

**DOI:** 10.64898/2026.03.23.713816

**Authors:** Hao Meng, Tobias Karmainski, Aziz Ben Ammar, Anka Sieberichs, Yannick Branson, Peer Vossen, Tobias Schwanemann, Hendrik Ballerstedt, Uwe T. Bornscheuer, Ren Wei, Lars M. Blank

## Abstract

Current mechanical and chemical recycling strategies address less than 10% of global plastic waste, necessitating alternative valorization routes. Biological upcycling via enzymatic depolymerization combined with microbial conversion of the resulting monomers offers a promising pathway to transform mixed plastic waste into valuable alternatives. Here, we employed a single engineered *Pseudomonas putida* KT2440 for simultaneous co-utilization of five plastic monomers including ethylene glycol, terephthalic acid, adipic acid, 1,4-butanediol, and L-lactic acid, which can be derived from enzymatic hydrolysis of polyethylene terephthalate (PET), polybutylene adipate-co-terephthalate (PBAT), polyester-polyurethanes (PUs), and polylactic acid (PLA). Continuous fermentation over 21 days with alternating mixed-monomer feeds achieved steady state growth and complete substrate depletion, yielding adaptive mutations that informed iterative strain improvement. Further engineering enabled the biosynthesis of (*R*)-3-hydroxybutyrate (R-3HB), and 0.70 g L^−1^ R-3HB was produced directly from enzymatic hydrolysates of blended PET, PBAT, and TPU. These results establish a viable bio-based approach for upcycling realistic mixed plastics into value-added bioproducts.

## Introduction

Plastics are indispensable to modern society, and annual production has increased sharply since 1950, reaching ∼430 Mt by 2024 [1]. Less than 10% of plastic waste is recycled; most is landfilled, incinerated, retained in use, or mismanaged [2], resulting in pervasive environmental leakage and associated ecosystem and human-health impacts [3,4]. Transitioning to a circular plastic economy will therefore require continued advances in both closed-loop and open-loop recycling technologies [5,6], with biotechnology emerging as a viable option to support this shift [7,8].

Enzymatic depolymerization of plastic waste into monomers or smaller molecules represents a promising and sustainable recycling strategy, with significant progress achieved in recent years [9–11]. This strategy is most effective for hydrolysable polymers containing backbone heteroatoms, including ester bonds in polyethylene terephthalate (PET), poly(butylene adipate-terephthalate) (PBAT), and polylactic acid (PLA); urethane bonds in polyurethanes (PUs); and amide bonds in polyamides (PAs) [12–14]. Conversely, the enzymatic degradation of non-hydrolysable plastics with C-C backbones (e.g., polyolefins or polystyrene) remains controversial, and especially the reported single-enzyme systems are questionable [15–17]. Instead, viable conversion of non-hydrolysable plastics typically requires a combinatorial approach that integrates chemical oxidation with multi-enzyme catalysis to produce intermediates for open-loop upcycling instead of closed-loop recycling [18–20].

PET is the industrial benchmark for plastic recycling [1] and has spearheaded enzymatic depolymerization research for over two decades [21,22]. Following years of extensive studies, numerous α/β-hydrolase fold superfamily enzymes have been discovered and engineered to rapidly depolymerize pretreated PET waste [21–23]. Notable benchmarks include variants of *Ideonella sakaiensis* PETase (*Is*PETase) and leaf-branch compost cutinase (LCC) [24–27], which have recently enabled the industrial demonstration of enzymatic PET recycling to achieve Technology Readiness Level (TRL) 8 (https://www.carbios.com/en/carbios-obtains-building-and-operating-permits/). Beyond PET, novel hydrolases capable of cleaving amide and urethane bonds in PAs and PUs, respectively, have also been identified [28–31]. These advances facilitate mixed-plastic depolymerization using mixed enzymes [19]. For example, Branson *et al*. demonstrated a one-pot dual-enzyme system that effectively depolymerizes PET, PBAT, and a thermoplastic polyester–polyurethane (TPU) [32].

Beyond closed-loop monomer recovery, these depolymerization products can serve as microbial feedstocks, providing carbon and energy for the biosynthetic production of value-added compounds via open-loop upcycling [10,33]. Several bacteria have been identified or engineered to utilize plastic monomers, including *Escherichia coli* [34,35], *Ideonella sakaiensis* [25], *Pseudomonas umsongensis* [36], *Acinetobacter baylyi*[37], *Paracoccus denitrificans* [38], *Pseudomonas taiwanensis* [39,40], and *Rhodococcus jostii* [41,42]. Among these, *Pseudomonas putida* KT2440 represents a particularly attractive catabolic platform due to its versatile metabolism, high stress tolerance [43–45], and a sophisticated genetic toolkit [46–49]. However, to date, its metabolic capacity has predominantly been engineered to utilize individual monomers [50–52] or monomers derived from a single plastic type [53,54]. For more complex mixed plastics, engineered microbial consortia have been explored, although their broader application remains limited [55–57].

To address the high complexity encountered under realistic plastic waste conditions, the metabolic capability of a single strain must be expanded. In this study, *P. putida* was selected as the host, and multiple plastic monomer metabolic pathways were consolidated into a single strain via chromosomal integration. The resultant “ETAB” strain was subsequently evaluated on mixed monomers across multiple cultivation scales, including continuous fermentation, where robust performance was observed. Based on genome re-sequencing of high-performing mutants isolated during continuous cultivation, reverse-engineering strategies were applied to further optimize the strain, which was additionally modified for (*R*)-3-hydroxybutyrate (R-3HB) production. Moreover, the efficiency in R-3HB production of the engineered strain was validated on mixed monomers derived from enzymatically depolymerized mixed plastics.

## Results

### Designing a single *P. putida* strain for the co-metabolization of EG, TA, AA, and BDO

The individual microbial utilization of the arguably four most common hydrolysable plastic-derived monomers, ethylene glycol (EG), terephthalic acid (TA), adipic acid (AA), and 1,4-butanediol (BDO), has been extensively studied (Figure 1A) [36,50–52,58]. *P. putida* natively harbors the genes encoding catabolism pathways for EG and BDO [51,59]. However, EG assimilation requires the activation of the *gcl* operon (*gcl*:*hyi*:*glxR*:*ttuD*:*pykF*) through deletion of the *gclR* repressor [50], and overexpression of the *glc* operon (*glcDEFG*) has been shown to further enhance growth [53,58,60]. BDO can be slowly assimilated by wild-type *P. putida*, but its utilization improves when PP_2046 is replaced with a constitutive P*_14g_* promoter [51].

**Figure 1.**
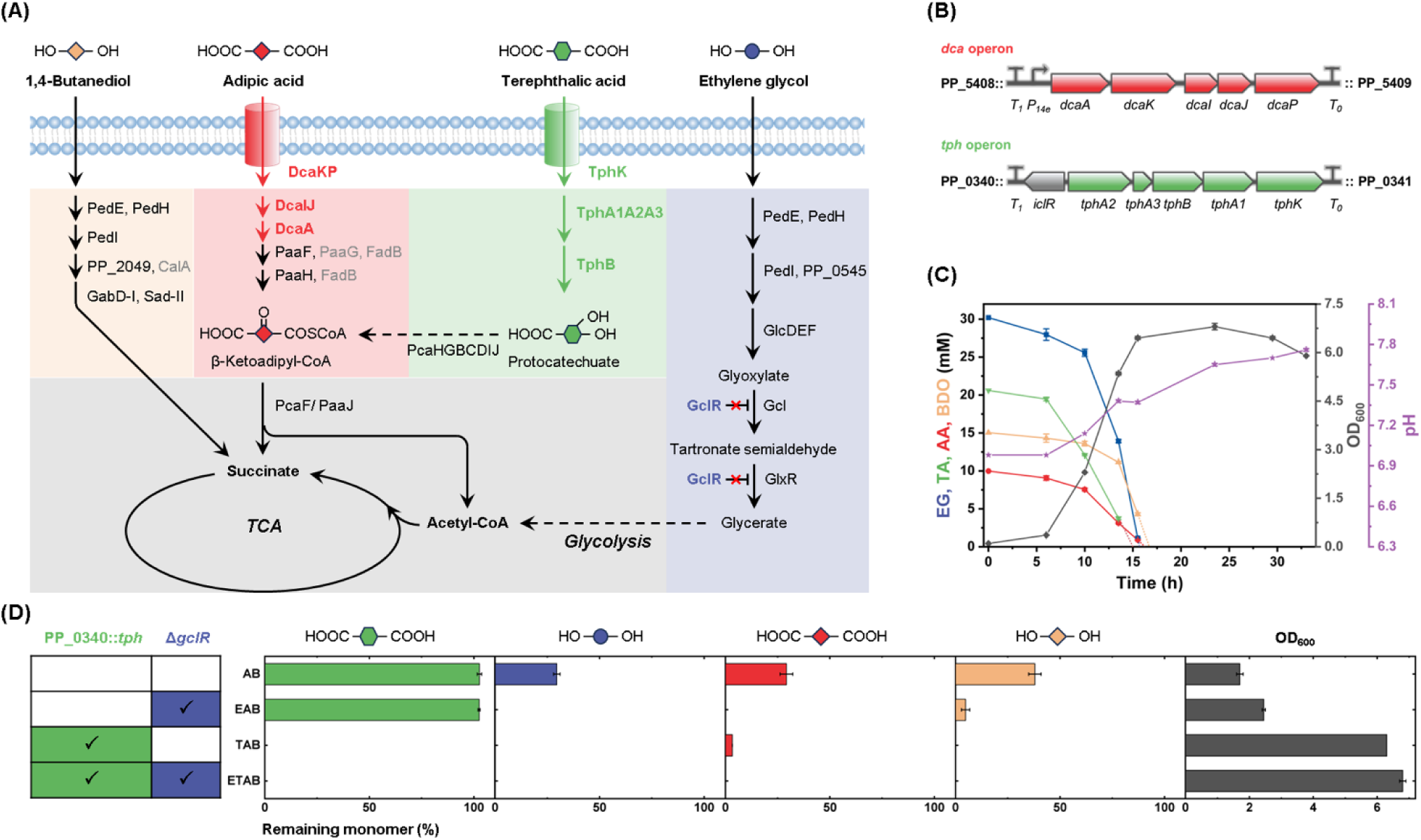
Integration of four metabolic pathways into a single *P. putida* strain and its functional validation. (A) Metabolic pathways for the assimilation of EG, TA, AA, and BDO. Symbols indicate structural units: rhombus, C4 unit “(–CH_2_–)_4_”; hexagon, benzene ring; circle, C2 unit “(–CH_2_–)_2_”. (B) Genetic organization of the integrated *tph* operon (*iclR*-*tphA2A3BA1K*) from *Pseudomonas umsongensis* GO16 for TA assimilation, and the *dca* operon (*P_14e_-dcaAKIJP*) from *Acinetobacter baylyi* for AA assimilation. (C) Time course of the concentrations of EG, TA, AA, and BDO in culture supernatants, together with growth and pH profiles of ETAB during the 33-hour cultivation on mixed monomers. (D) Remaining monomer percentages for strains AB, EAB, TAB, and ETAB, along with corresponding OD_600_ values after 24 h of cultivation. Error bars represent standard deviations among biological duplicates.

In contrast, *P. putida* lacks both TA and AA transport systems, as well as the enzymatic pathways required to convert TA to protocatechuate (PCA) and AA to 2,3-didehydroadipyl-CoA, necessitating the introduction of heterologous pathways. Previously, Ackermann *et al*. introduced the *dca* operon (*dcaAKIJP*) from *Acinetobacter baylyi* into *P. putida*, along with additional genomic modifications, enabling efficient AA utilization [52]. Several bacteria, including *Comamonas* sp., *Rhodococcus jostii*, *P. umsongensis*, and *I. sakaiensis*, harbor the missing enzymatic steps for TA metabolism [25,36,61,62]. Early engineering efforts confirmed that the combination of *tphK* from *R. jostii* with one *tph* operon from *Comamonas* sp. showed superior performance [53]. More recently, a compact *tph* operon (*iclR*-*tphA2A3BA1K*) was identified in *P. umsongensis*, in which the MFS transporter *tphK* and the *tphA2A3BA1* genes are co-localized within a single transcriptional unit, representing an efficient configuration for TA metabolism [36].

To develop a single *P. putida* strain capable of utilizing all four monomers, we hypothesized that this could be achieved by selectively combining the modifications described above. We started from the strain KT2440ge ΔP*_paaF_*-*paaYX*::P*_14g_* Δ*psrA* (referred to as AB in this study), obtained from the Wierckx group, which already carries the *dca* operon from *Acinetobacter baylyi* (Figure 1B) together with additional modifications enabling AA utilization [52]. Two more modifications were then introduced. First, to enable TA metabolism, the *tph* operon from *P. umsongensis* GO16 was integrated at the PP_0340 locus (Figure 1B) [63]. Second, *gclR* was deleted to activate EG assimilation (Figure 1A). By integrating these modifications, three engineered strains, EAB, TAB, and ETAB, were constructed (Figure 1D).

These four strains were subsequently evaluated for growth on mixed monomers (Figure 1C, D, Figure S1). As expected, the parental AB strain and the EAB strain did not consume TA due to the absence of the *tph* operon. Following the deletion of *gclR*, EAB consumed other monomers more rapidly, likely because EG assimilation was activated, thus contributing to biomass formation. However, after 24 h, EAB cultures exhibited significant cell clumping for reasons that remain unclear. Moreover, the final pH of the AB and EAB cultures decreased relative to the initial value, suggesting the accumulation of unidentified acidic byproducts. In contrast, both TAB and ETAB efficiently co-consumed all four monomers, achieving complete depletion within 17 h and reaching similar final pH values (∼7.8). Notably, ETAB displayed improved growth performance compared with TAB, reaching an approximately 8% higher maximum OD_600_. This improvement is likely attributable to the activated EG assimilation. Residual monomer concentrations at 24 h and the maximum OD_600_ values are summarized in Figure 1D for direct comparison. In summary, an engineered *P. putida* strain designated ETAB was obtained, exhibiting robust growth on mixed monomers.

### Physiological characterization of the ETAB strain

To further evaluate the ETAB strain performance on mixed monomers, monomer preference was assessed. The ETAB strain was cultivated on five individual monomers, EG, TA, AA, BDO, and L-lactic acid (LA), as sole carbon sources (Figure 2A). Substrate utilization was compared by online monitoring of the oxygen transfer rate (OTR), which serves as a proxy for the cells’ respiratory activity and substrate consumption. The results indicate a clear preference order, with LA being most rapidly utilized, followed by TA, AA, BDO, and EG. As PET and PUs are among the most widely used hydrolysable plastics, their corresponding monomers were selected for subsequent experiments. PET consists of TA and EG in a 50:50 molar ratio, whereas the soft segments of PUs can contain EG, AA, and BDO in a 25:50:25 molar ratio [57] (Figure 2B).

**Figure 2.**
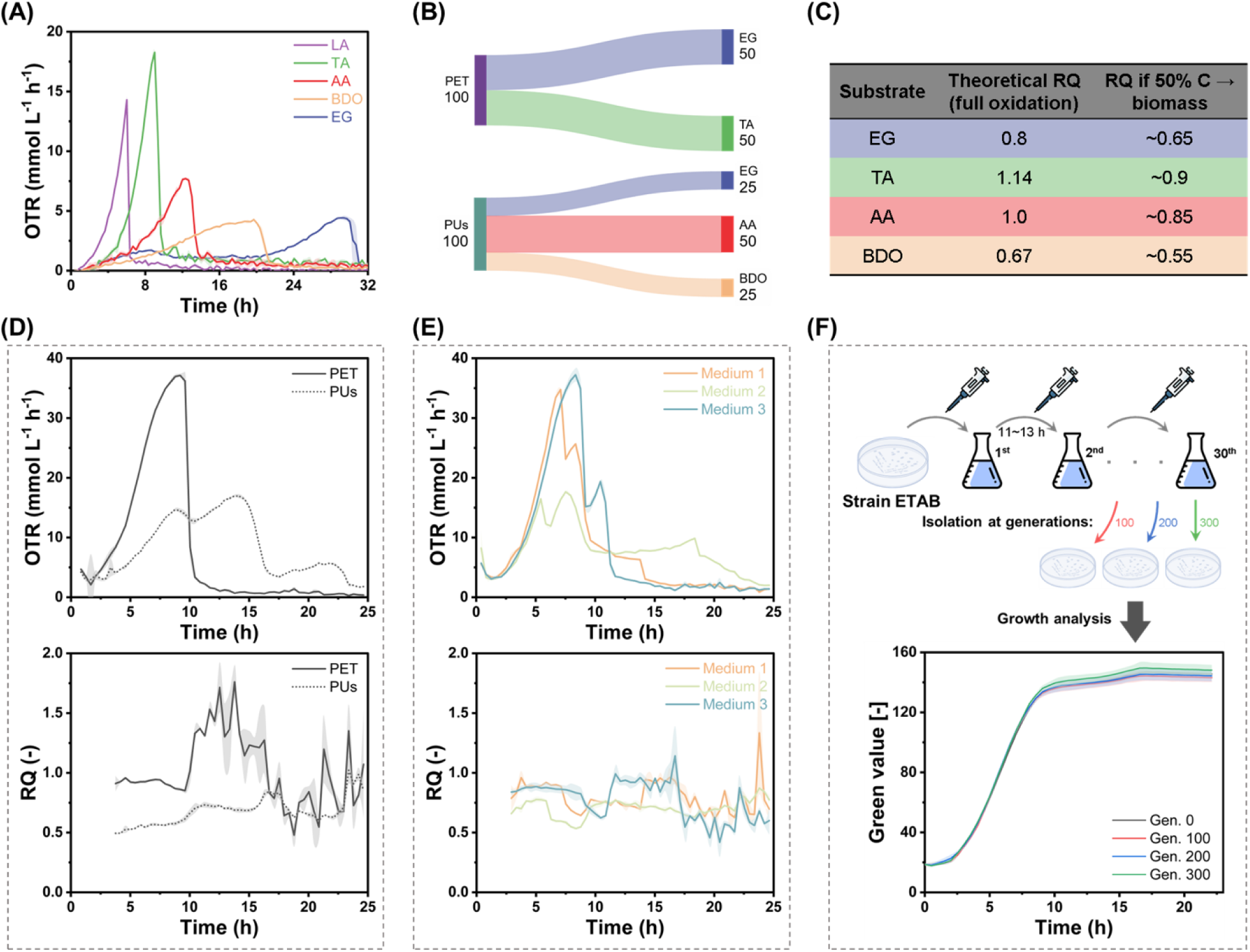
Physiological characterization of the ETAB strain. (A) Comparison of growth on individual monomers by online monitoring of oxygen transfer rate (OTR) in a 96-deepwell plate with a Kuhner microTOM shaker. Substrate concentrations: EG, 30 mM; TA, 10 mM; AA, 10 mM; BDO, 15 mM; LA, 20 mM. (B) Metabolizable monomer composition of PET and soft segments of PUs. (C) Theoretical respiratory quotient (RQ) values of the individual monomers. (D) OTR and RQ profiles of ETAB cultivated on single plastic (PET or soft segments of PUs) monomers at a total monomer concentration of 60 mM. (E) OTR and RQ profiles of ETAB cultivated on mediums 1, 2 and 3. (F) Evaluation of genetic stability of the ETAB strain during serial cultivation on LB medium. After 100, 200, and 300 generations, cultures were isolated on LB agar plates, and six single colonies from each time point were tested for growth on medium 1. Shaded areas represent standard deviations of biological triplicates (n=3).

The ETAB strain was subsequently evaluated on single plastic (PET or soft segments of PUs) monomers, with substrate utilization monitored by OTR and respiratory quotient (RQ), the latter indicating CO_2_-to-O_2_ stoichiometry and thus changes in the dominant catabolic substrate (Figure 2D). When cultivated on PET monomers, the OTR profile exhibited a single peak, indicating simultaneous co-consumption of TA and EG. In contrast, cultivation on monomers from soft segments of Pus resulted in three closely spaced OTR peaks. Based on the theoretical RQ values of the individual monomers (Figure 2C), the measured RQ profiles suggest sequential substrate utilization with AA clearly not utilized initially. This pattern differs from the preference observed during growth on individual monomers. These results indicate that the hierarchy of monomer utilization can shift under mixed-substrate conditions.

Online OTR and RQ measurements were further used to compare ETAB performance on three different monomer compositions (based on fluctuating PET and soft segments of PUs ratios, at constant total monomer concentration of 60 mM), reflecting variances in plastic waste: Medium 1. 22.5 mM EG, 15 mM TA, 15 mM AA, and 7.5 mM BDO; Medium 2. 27 mM EG, 24 mM TA, 6 mM AA, and 3 mM BDO; and Medium 3. 18 mM EG, 6 mM TA, 24 mM AA, and 12 mM BDO (Figure 2B, E). Offline measurements of OD_600_ and residual monomer concentrations were conducted as references (Figure S2). With decreasing proportions of PET monomers, overall growth performance declined. Notably, using medium 3, EG, TA, and BDO were depleted within 10 h, whereas complete AA consumption took 24 h, indicating that AA metabolism is strongly affected by mixed-substrate conditions. The maximum specific growth rates (µ_max_) were 0.51, 0.50, and 0.41 h^−1^ on medium 1, 2 and 3, respectively, showing a decrease with lower PET monomer content. The different effects resulting from the applied monomer composition shaped the design of subsequent continuous fermentation experiments.

To assess the genetic stability of ETAB, the strain was serially cultivated on LB medium for approximately 300 generations, with samples collected every 100 generations. Isolates were obtained on LB agar plates, and six randomly selected colonies from each time point were evaluated on mixed monomers. As shown in Figure 2F, strains from all generations exhibited comparable performance, indicating stable phenotypes during prolonged cultivation on LB medium.

### Continuous fermentation using alternating mixed-monomer feeds

To evaluate its growth and metabolic performance under realistic plastic recycling conditions, the engineered ETAB strain was cultivated for 21 days in a continuous fermentation with alternating mixed-monomer feeds (Figure 3A). The process was initiated with a 9 h-batch phase using medium 1, during which TA was depleted first, followed by EG and BDO, resulting in a CDW of 2.8 g L^−1^ (Figure 3B). Continuous operation was then started at a dilution rate of 0.1 h^−1^.

**Figure 3.**
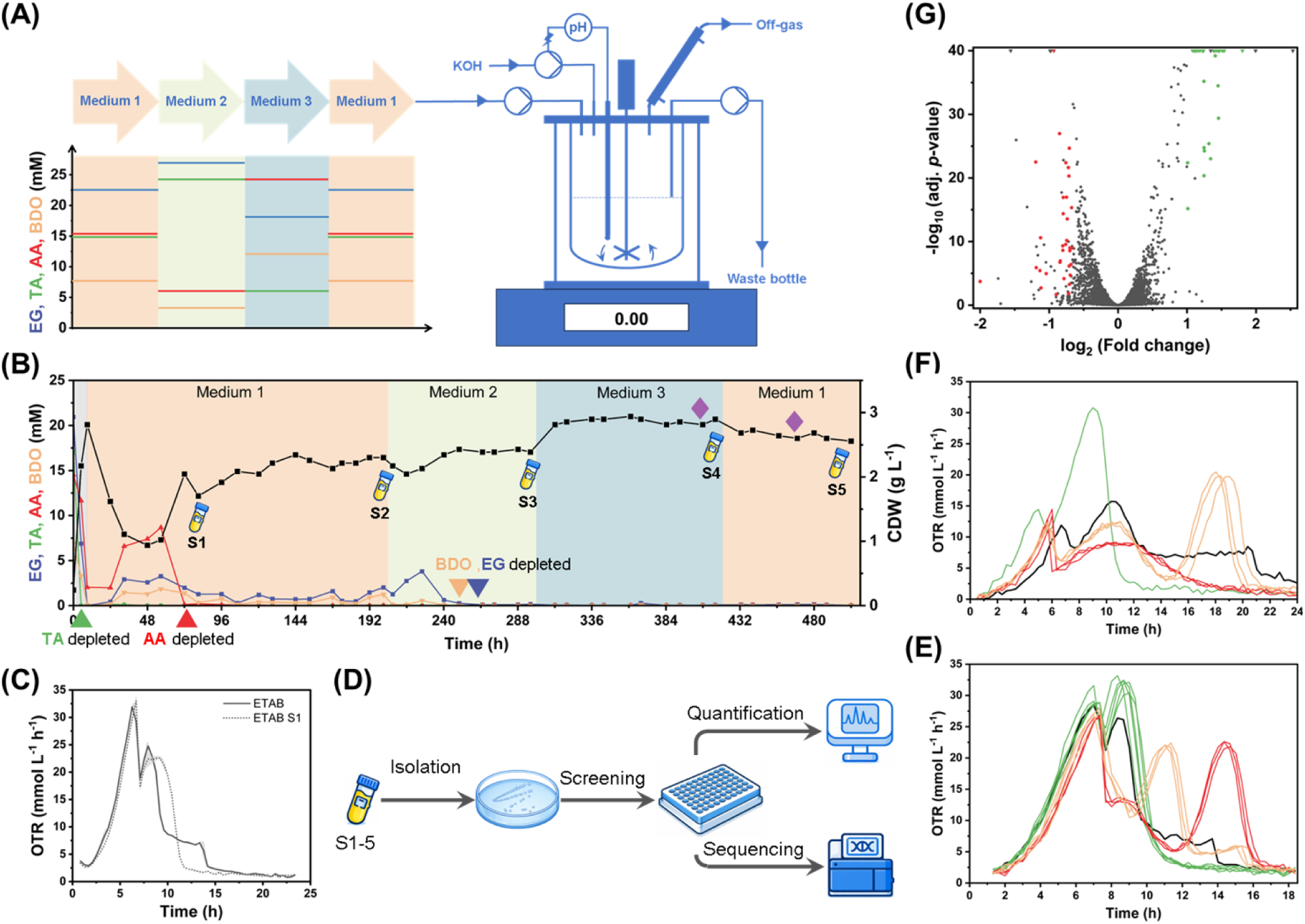
In-depth analysis of continuous fermentation with the ETAB strain. (A) Overview of the fermentation setup and the alternating mixed-monomer feeds. (B) Biomass formation and consumption of the four monomers, EG, AA, TA, and BDO, during the 21-day continuous fermentation. Colored triangles indicate depletion of individual monomers. Blue sample tube icons labeled S1-S5 denote the five sampling time points for strain isolation, and purple diamonds indicate the two sampling points used for transcriptome analysis. (C) Comparison of the ETAB strain and the strain mixture from S1 on medium 1, monitored online via the OTR. Shaded areas represent standard deviations from biological triplicates (n=2). (D) Overview of the workflow from isolating the 5 rounds of strain sampling with the following screening by monitoring OTR on mixed monomers, quantification by HPLC for the residue monomers and genome resequencing. (E) Monitored OTR of representative isolates from S1 on medium 1. (F) Monitored OTR of representative isolates from S4 on medium 3. (G) Overview of the transcriptome analysis comparing the medium 3 phase with the following medium 1 phase. Red and green symbols indicate significantly downregulated and upregulated known genes, respectively.

During the initial continuous phase (medium 1), biomass declined rapidly to 0.9 g L^−1^, accompanied by marked AA accumulation (up to 58%), while EG and BDO also accumulated. Notably, AA was subsequently depleted within 15 h, and biomass recovered to 2.0 g L^−1^. Cells collected at this time point (S1) exhibited significantly improved substrate utilization compared with the parental ETAB strain, as revealed by OTR analysis (Figure 3C), suggesting adaptive mutations that enhanced AA metabolism under mixed-substrate conditions.

As cultivation proceeded through successive feeding phases, the system reached stable steady states at each phase. Early phase transitions were accompanied by transient biomass decreases; however, later switches led to immediate recovery to steady-state levels without observable lag, indicating increased regulatory flexibility and adaptation to fluctuating monomer compositions. In the latter half of the fermentation, all monomers were completely consumed. The fermentation was terminated at 504 h, with off-gas profiles (OTR, CTR, and RQ) shown in Figure S3A.

Overall, the long-term continuous cultivation not only demonstrated robust utilization of mixed monomers but also drove adaptive improvements that enhanced metabolic performance under dynamically changing substrate conditions. During the process, a maximum total monomer uptake rate of 4.38 mmol g^−1^ h^−1^ and a maximum biomass yield of 0.45 g g^−1^ were achieved (Figure S3B), with potential for further enhancement at higher dilution rates. To our knowledge, this study is the first to report a continuous cultivation process that achieves stable steady states under alternating mixed-monomer compositions with complete depletion of all supplied monomers [64].

Cells collected at five time points (S1–S5) were isolated on agar plates, and 94 colonies from each population were randomly selected for OTR-based performance screening in comparison with the parental ETAB strain (Figure 3D). Among the S1 isolates, five strains exhibited improved performance (in green) relative to the parental strain (black) (Figure 3E). In contrast, only one improved isolate was identified from the S4 population (Figure 3F). No additional significantly enhanced strains were detected among the total 470 isolates screened. Notably, some isolates displayed reduced performance compared with the parental strain. This may reflect the relatively moderate selective pressure imposed by the fixed dilution rate (0.1 h^−1^), which could have allowed the emergence of less competitive phenotypes.

Samples for transcriptome analysis were collected during the final two feeding phases of the continuous fermentation (purple diamonds, Figure 3B), and the overall results are summarized in Figure 3G. Upon switching between these phases, TA concentration markedly increased (1.5-fold). Correspondingly, transcriptomic analysis revealed an approximately 1.7-fold upregulation of the *tph* operon (*tphA2A3BA1K*), indicating that their expression responds to TA concentration. In addition, the *alg* operon involved in alginate biosynthesis was also upregulated (on average ∼1.7-fold), which may reflect increased cellular stress associated with elevated TA levels [65].

### Genome re-sequencing and reverse engineering to further improve the ETAB strain

The genomes of selected isolates from the continuous fermentation were re-sequenced, revealing two major point mutations in PP_2046 (identified in strain S1-13) and *pcaR* (identified in strain S3-86) (Figure 4A). Reverse engineering was performed by introducing these mutations individually and in combination into the parental ETAB background, generating three derivative strains: ETAB V2 carrying PP_2046^c.-22T>A^; ETAB V3 carrying *pcaR*^D129Y^; and ETAB V4 harboring both mutations.

**Figure 4.**
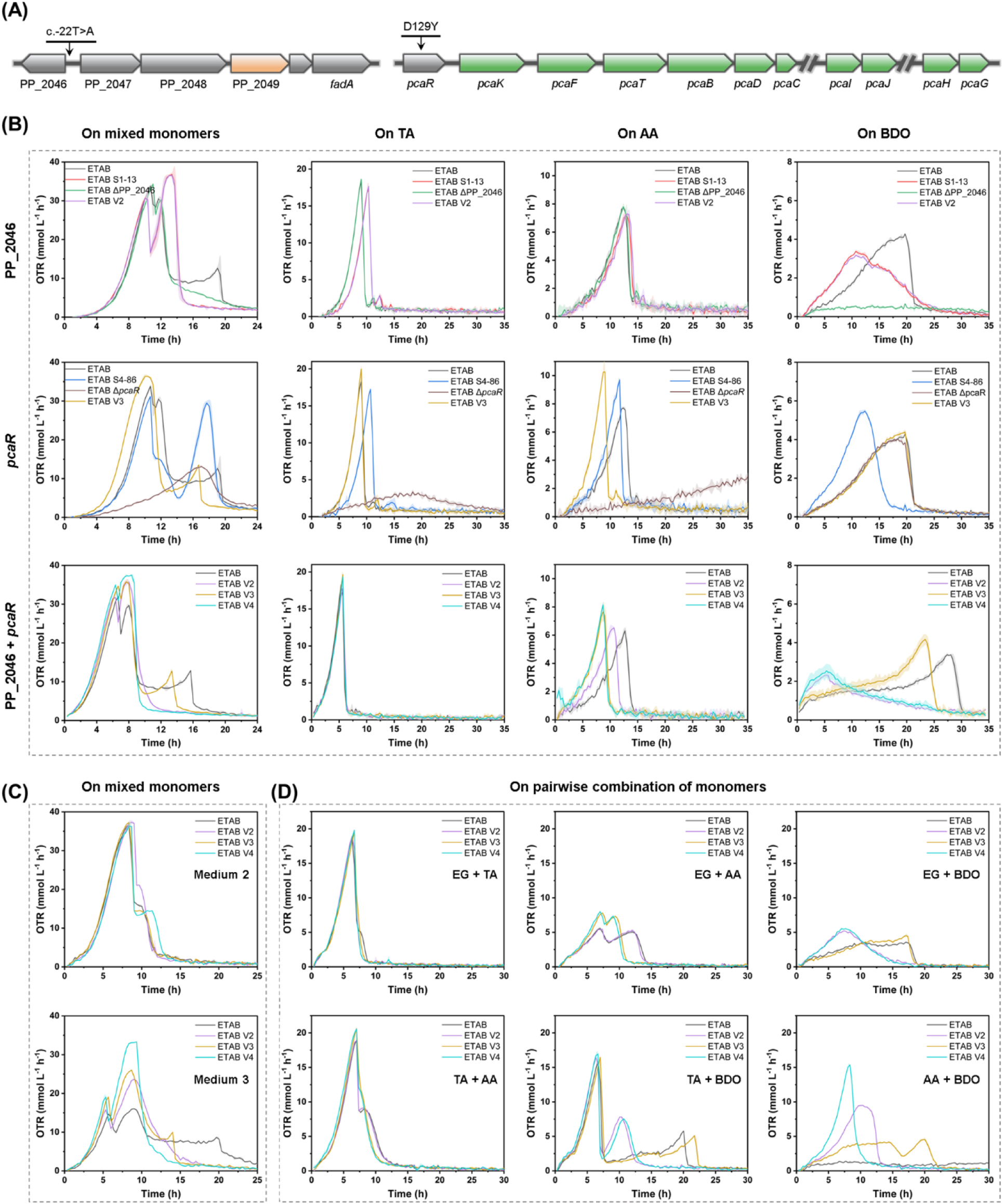
Reverse engineering of mutations identified from continuous fermentation. (A) Two point mutations revealed by genome re-sequencing. (B) Comparison of reverse-engineered strains with evolved, parental, and respective deletion strains based on substrate utilization. Strains were cultivated on medium 1, or on single monomer with concentrations of 10 mM TA; 10 mM AA, and 15 mM BDO. Glucose was used as the preculture carbon source when compared with the deletion strains, whereas mixed-monomer medium was used for other pre-cultivations. (C) Comparison of reverse-engineered strains with the parental strain on mediums 2 and 3. (D) Comparison of reverse-engineered strains with the parental strain on pairwise combinations of monomers (10 mM each). Shaded areas represent standard deviations from biological triplicates (n=3).

These strains were subsequently compared by monitoring the OTR with the parental and reference strains under different cultivation conditions (Figure 4B-D, Figure S5). Regarding the PP_2046 mutation, ETAB V2 exhibited improved performance on BDO, pairwise substrate combinations, and mixed monomers, particularly on the AA + BDO combination. In contrast, growth on TA, AA, and their combination was slightly reduced. The phenotype closely resembled that of strain S1-13 and the strain mixture ETAB S1 shown in Figure 3C, confirming successful reconstruction of the evolved metabolic characteristics. Comparison between ETAB and ETAB ΔPP_2046 further demonstrated that deletion of this regulator affects BDO metabolism (Figure 4A). For the *pcaR* mutation, ETAB V3 displayed enhanced utilization of both TA and AA, resulting in significantly faster growth on mixed monomers. Deletion of *pcaR* impaired TA metabolism and led to accumulation of protocatechuate (PCA) (Figure S4B); unexpectedly, it also abolished AA consumption, indicating a previously unrecognized dependence of AA metabolism on *pcaR*. Notably, the ETAB V4 strain demonstrated improved growth across nearly all tested conditions, including the four-monomer mixture, pairwise combinations, and individual substrates (Figure 4B-D, Figure S5), thereby expanding the design space for rational engineering towards plastic monomer utilization.

### Further engineering for R-3HB production from mixed plastic monomers

Following the efficient utilization of mixed monomers, we next focused on the production of (*R*)-3-hydroxybutyrate (R-3HB), the monomer of the biodegradable polymer poly(3-hydroxybutyrate) (PHB). Beyond its role as a bioplastic precursor, R-3HB can function as a natural ketone body in humans, with reported applications in cancer metabolism studies and as a clinical biomarker [66,67].

It also serves as a valuable chiral building block in chemical synthesis. Recently, R-3HB production titers of up to 105 g L^−1^ have been achieved in an engineered *E. coli* strain [68], demonstrating the feasibility of large-scale bioproduction. However, to date, R-3HB production in *P. putida* has not been reported.

For R-3HB production (Figure 5A), the *phaA3* and *phaB* genes encoding β-ketothiolase and NADPH-dependent acetoacetyl-CoA reductase, respectively, were introduced from *Pseudomonas pseudoalcaligenes* CECT5344 [69]. These genes were co-expressed in *P. putida* with the native *tesB* under the control of the strong constitutive promoter P*_14ffg_* (Figure 5B) [70]. However, no extracellular R-3HB was detected when the resulting KT2440-PP strain was cultivated on glucose, consistent with a recent report [71]. To optimize the module, deletion candidates were evaluated in the KT2440-PP background (Figure S6). Deletion of *hbdH* and *atoAB* was required for R-3HB accumulation, whereas additional deletion of *hbd* further minimized competing flux and increased the R-3HB titer by ∼12%. Based on these results, the R-3HB production module was finalized (Figure 5A).

**Figure 5.**
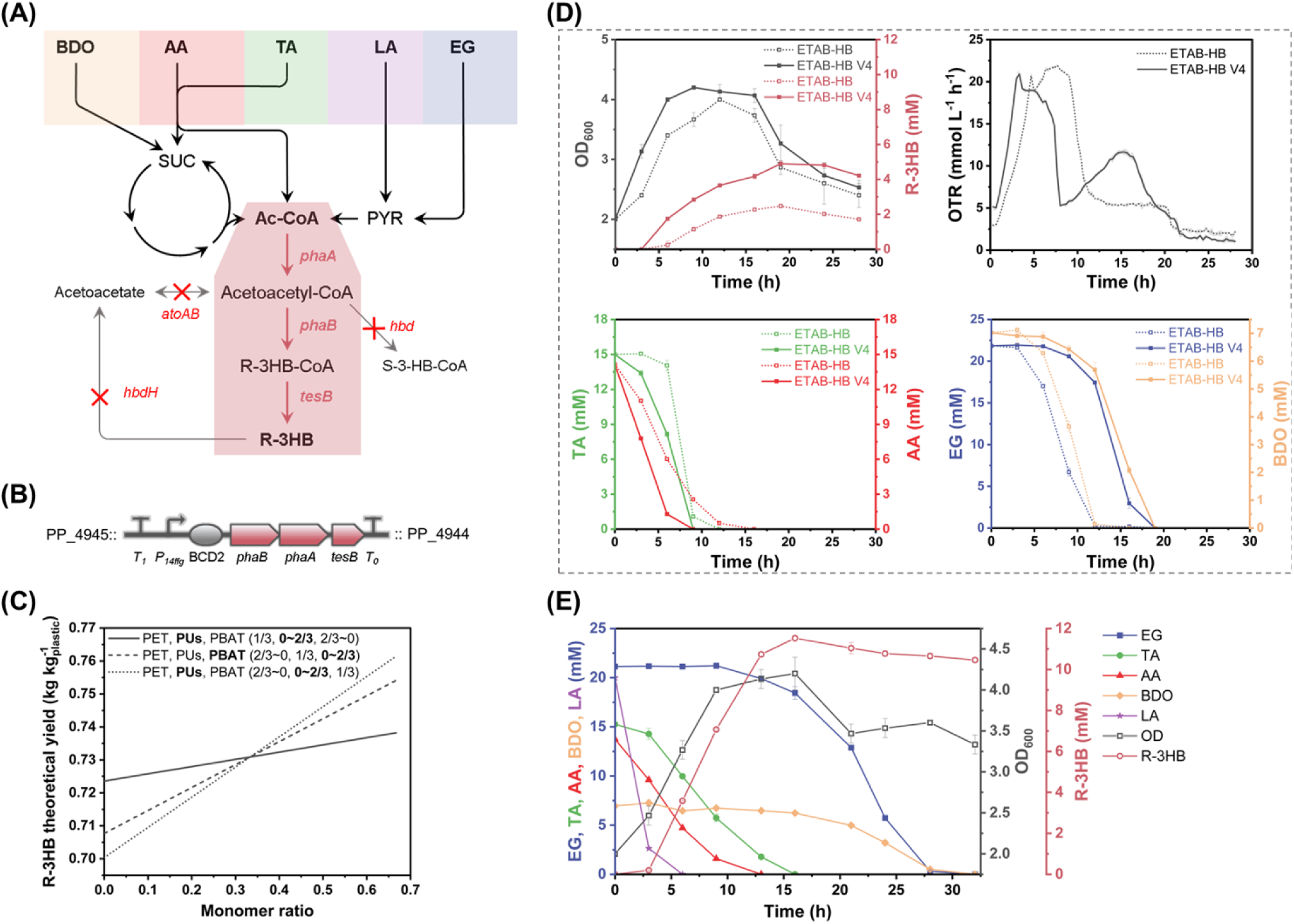
R-3HB production from mixed plastic monomers. (A) Overview of the assimilation pathways for the five plastic monomers, EG, TA, AA, BDO, and LA, and the corresponding genetic modifications introduced for the R-3HB production module. (B) Genomic integration of the operon (P*_14ffg_*-BCD2-*phaBA3*-*tesB*) for R-3HB production. (C) Theoretical yield (kg kg^−1^) of R-3HB from PET, PUs, and PBAT monomer mixtures at different ratios. (D) Comparison of R-3HB producer strains ETAB-HB and ETAB-HB V4 with respect to R-3HB production, growth, and monomer consumption during cultivation on medium 1 (NH_4_^+^ concentration 3 mM) in shake flasks, together with online monitoring of the oxygen transfer rate (OTR) during the same cultivation in 96-deepwell plates. (E) R-3HB production, growth, and monomer consumption by strain ETAB-HB V4 during cultivation on medium 1 (with an additional 20 mM LA; NH_4_^+^ concentration 3 mM) in shake flasks. Error bars represent standard deviations among biological triplicates (n=3).

In parallel, a metabolic model based on iJN1463 [72] was expanded to incorporate the TA and AA assimilation pathways. Based on this model, the theoretical yields of R-3HB from mixed plastics (PET, PUs, and PBAT) at different ratios were calculated and predicted to exceed those obtained from glucose (0.58 kg_R-3HB_ kg_Glucose_^−1^), highlighting the potential of plastic-derived substrates (Figure 5C).

This R-3HB production module was then integrated into the ETAB and ETAB V4 backgrounds, generating the strains ETAB-HB and ETAB-HB V4, which were subsequently evaluated for R-3HB production on mixed monomers under nitrogen-limited conditions (Figure 5D). Although both strains required a similar overall time to deplete all monomers, ETAB-HB V4 achieved approximately double the R-3HB titer (4.9 mM, 0.51 g L^−1^, 0.04 g_R-3HB_ g_monomers_^−1^) compared with ETAB-HB (2.5 mM, 0.26 g L^−1^, 0.08 g_R-3HB_ g_monomers_^−1^). The primary phenotypic difference between the two strains was that ETAB-HB V4 consumed TA and AA more rapidly than ETAB-HB, accompanied by faster biomass accumulation and earlier nitrogen depletion, whereas EG and BDO were utilized later. The mechanistic basis for this altered substrate distribution and improved R-3HB production remains unclear.

The ETAB-HB V4 strain was subsequently cultivated in a medium supplemented with an additional 20 mM LA (Figure 5E). LA can be readily converted into the R-3HB precursor acetyl-CoA via a short, two-step pathway (Figure 5A). Notably, supplementation with LA, which increased the total carbon input by only 21%, resulted in a 1.35-fold higher R-3HB titer (11.5 mM, 1.2 g L^−1^, 0.14 g_R-3HB_ g_monomers_^−1^). These results suggest that R-3HB production from the other four monomers is primarily limited by the efficiency of acetyl-CoA supply.

### Production of R-3HB from enzymatically depolymerized mixed plastics

For bio-based upcycling applications, mixed monomers undergoing microbial transformations are expected to originate from enzymatically depolymerized mixed plastics [7,10] (Figure 6A). To approximate post-consumer waste conditions, a polymer blend composed of PET, PBAT, and a thermoplastic polyester-polyurethane (TPU) was hydrolyzed using a sequential one-pot dual-enzyme system as described by Branson *et al.* [32]. The resulting hydrolysates were used directly to prepare the cultivation medium, in which the strain ETAB-HB V4 was cultivated. Comparable performance was observed relative to the cultivation on defined mixed monomers (Figure 6B). AA and TA were co-consumed first, accompanied by nitrogen depletion as indicated by OTR measurements (Figure 6C), followed by the co-consumption of EG and BDO. R-3HB production reached 6.6 mM (0.70 g L^−1^, 0.07 g_R-3HB_ g_monomers_^−1^) at 28 h, at which point all monomers were depleted. In addition, the TPU-derived diamine 4,4’-methylenedianiline (MDA) did not measurably impair growth at the detected concentration (0.12 mM), as previously reported [73]. Notably, the R-3HB titer continued to increase after complete substrate consumption, the underlying mechanism of which remains unclear. Overall, this experiment provides a proof-of-concept for the direct production of R-3HB from enzymatically hydrolyzed mixed plastics.

**Figure 6.**
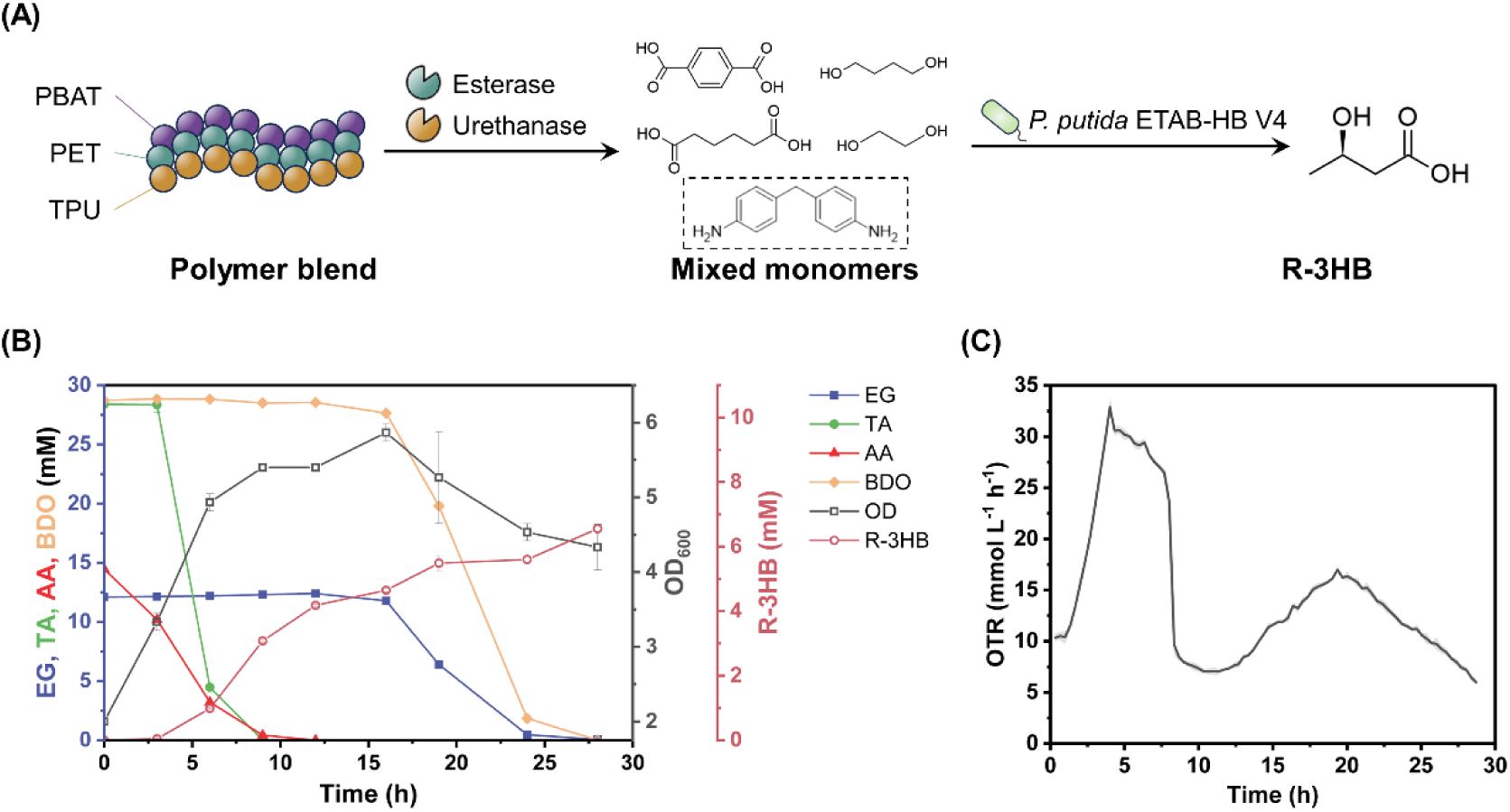
Production of R-3HB using monomers derived from enzymatically depolymerized blended polymers. (A) Workflow for R-3HB production from mixed plastics. (B) R-3HB production by strain ETAB-HB V4 in shake-flask cultures using media prepared from mixed plastic hydrolysates. (C) Online monitoring of the OTR during growth of strain ETAB-HB V4 on medium prepared from mixed plastic hydrolysates in 96-deepwell plates. Error bars represent standard deviations among biological triplicates.

## Discussion

In this study, we constructed a robust bacterial strain, *P. putida* ETAB, capable of simultaneously utilizing five plastic monomers, EG, TA, AA, BDO, and LA. Its performance on mixed substrates was validated through continuous fermentation with alternating mixed-monomer feeds, demonstrating stable and efficient utilization. Beneficial point mutations were identified during this process, enabling further strain improvement through reverse engineering. The strain was additionally engineered genomically to produce the versatile platform chemical R-3HB from mixed plastic hydrolysates. Together, these results highlight the utility of *Pseudomonas* metabolic flexibility for valorizing complex plastic-derived feedstocks, while underscoring the need to better channel carbon flux toward target products.

Online monitoring of respiratory activity (OTR, CTR, and RQ) provides a powerful alternative to conventional endpoint metabolite analysis, particularly for multi-substrate systems [74,75]. Respiration profiles resolved substrate utilization dynamics and revealed physiological constraints, including nitrogen and oxygen limitation, that were not readily captured by supernatant measurements alone. Moreover, respiration profiling under defined nitrogen conditions highlighted the critical role of nitrogen availability in mixed-monomer utilization. Although the reverse-engineered ETAB V4 strain achieved simultaneous co-consumption of mixed monomers (Figure 4B), preferential utilization of TA and AA rapidly depleted nitrogen, triggering a transient collapse in metabolic activity followed by recovery. This behavior suggests that efficient EG and BDO metabolism depends on sufficient nitrogen supply and illustrates the value of respiration-based monitoring for identifying hidden bottlenecks in mixed-substrate bioprocesses.

Substrate co-utilization in complex feedstocks is governed not only by pathway availability but by regulatory hierarchy. Here, we focused on the co-utilization of EG, TA, AA, and BDO, whereas LA and MDA was not analyzed in detail. Given the reported toxicity of MDA [73], future process designs may benefit from separating this compound during depolymerization/hydrolysate conditioning or replacing it with biologically less toxic diamines. As the most preferred carbon source (Figure 5E) [76], LA is rapidly converted to acetyl-CoA and did not interfere with other monomer assimilation (Figure S7), indicating that regulatory competition primarily occurs among the more complex aromatic and dicarboxylic substrates. Although *pcaF*-II was proposed to encode a functional 3-ketoadipyl-CoA thiolase activated via *psrA* deletion [52,77]; deletion of *pcaR* completely abolished AA metabolism despite activation of *pcaF*-II (Figure 4B), demonstrating a dominant role for *pcaF*-I. Consistently, the improved TA/AA co-metabolization observed in the *pcaR*^D129Y^ mutant likely reflects enhanced control of *pcaF-*I expression. Similarly, the PP_2046^c.-22T>A^ mutation markedly altered AA/BDO co-utilization (Figure 4D), revealing additional regulatory control points outside canonical degradation pathways. While the global regulator Crc has been implicated in mixed-substrate utilization [78,79] and intersects with PcaR-mediated regulation [77], our data demonstrate that broad regulatory deletions provide limited benefit (Figure S8) and may introduce unintended trade-offs, including unwanted accelerated product degradation. Collectively, these observations emphasize regulatory engineering as a key lever for optimizing mixed plastic monomer co-utilization.

As the extent of genome editing increases and heterologous pathways accumulate, strain robustness and metabolic burden become critical considerations. In this study, deletion of *gclR* or *psrA* primarily activated existing pathways [50,52], while the introduced *tph* operon retained native regulatory control and its expression was regulated by TA availability [36], minimizing unnecessary metabolic burden. In addition, three constitutive promoters (P*_14e_*, P*_14g,_* and P*_14ffg_*) were integrated into the genome to drive overexpression of the *dca* operon, *paaF*, and the R-3HB production module. Although strong constitutive promoters (P*_14e_*, P*_14g_*, and P*_14ffg_*) were integrated to drive key modules, yet no significant growth impairment was observed (Figure S9), and the genetic stability of ETAB was maintained over at least 300 generations in LB medium. Nevertheless, dynamic or self-regulated expression systems may further improve long-term performance and reduce metabolic load.

A remaining limitation is the relatively low carbon yield of R-3HB production from plastic monomers, likely attributable to inefficient conversion of substrates into acetyl-CoA. EG assimilation via the glyoxylate carboligase pathway entails 50% carbon loss, while TA, AA, and BDO metabolism involves carbon loss during conversion from malate to acetyl-CoA. Potential solutions include alternative EG routes (e.g., acetyl-phosphate synthase or diol dehydratase pathways) to eliminate carbon loss [80–82] and engineering of a reverse glyoxylate shunt to enable theoretical carbon conservation toward acetyl-CoA [83,84]. Realizing these gains will require enzyme optimization and pathway balancing under relevant process conditions.

### Concluding remarks

In summary, this work demonstrates a microbial route for converting enzymatically depolymerized mixed plastic monomers into R-3HB using a single engineered *P. putida* strain, while also revealing regulatory and process constraints that shape mixed-substrate performance. Despite tolerating considerable substrate complexity, the system remains distant from the heterogeneity encountered in real waste streams, including the enormous diversity of additives [85], and the metabolic capacity of a single strain will ultimately reach its limits. Future recycling strategies may therefore benefit from cascade configurations (e.g., sequential reactors) or rationally designed consortia to distribute metabolic functions and burdens across specialized strains. Notably, the ETAB strains were constructed as fully genomically engineered, marker-free systems with constitutive or self-induced expression architectures that avoid external inducers, providing a broadly applicable design framework for environmentally relevant biotechnological applications. Overall, these findings lower barriers to upcycling complex plastic waste streams and advance open-loop recycling strategies that convert plastic waste into value-added products.

## EXPERIMENTAL MODEL AND STUDY PARTICIPANT DETAILS

### Bacterial strains, medium composition, and culture conditions

All the bacterial strains and plasmids used in this study are listed in Tables S1 and S2, respectively. *E. coli* PIR2 was used as a cloning host and also as a donor strain during parental mating. *E. coli* HB101 strain was used as a helper strain for the parental mating. Lysogeny broth (LB) (containing 10 g L^−1^ tryptone, 5 g L^−1^ yeast extract, and 5 g L^−1^ NaCl) and mineral salts medium (MSM) [86] were used for all cultivations. The MSM contained 5 g L^−1^ glucose or different mixed plastic monomers as carbon sources, (NH_4_)_2_SO_4_ as the nitrogen source, and sodium/potassium phosphate as the buffering system with an initial pH of 7.0. The medium was supplemented with 1× trace element solution containing (per liter): 10 mg EDTA, 0.1 mg MgCl_2_·6H_2_O, 2 mg ZnSO_4_·7H_2_O, 1 mg CaCl_2_·2H_2_O, 5 mg FeSO_4_·7H_2_O, 0.2 mg Na_2_MoO_4_·2H_2_O, 0.2 mg CuSO_4_·5H_2_O, 0.4 mg CoCl_2_·6H_2_O, and 1 mg MnCl_2_·2H_2_O.

In this study, 1× phosphate buffer was defined as 3.88 g L^−1^ K_2_HPO_4_ and 1.63 g L^−1^ NaH_2_PO_4_ (36 mM total phosphate), and 1×nitrogen source corresponded to 2 g L^−1^ (NH_4_)_2_SO_4_. For shake-flask and microtiter plate cultivation, 3× buffer was used together with 1× (NH_4_)_2_SO_4_, 1× trace elements. For the continuous fermentation, 1× buffer was used together with 1.5× (NH_4_)_2_SO_4_, 2× trace elements. For R-3HB production experiments, a 4×buffer system (composed of 3× PIPES and 1× phosphate) was used with an initial pH of 6.4, and 0.2× (NH_4_)_2_SO_4_ was applied to establish nitrogen-limited conditions (0.25× (NH_4_)_2_SO_4_ was used for mixed plastic hydrolysates).

For the cultivation, strains were first streaked from glycerol stocks stored at −80 ℃ onto LB agar plates and incubated overnight. The strains were then used to inoculate liquid LB medium, which were used as first pre-cultures. These were subsequently transferred into MSM supplemented with glucose (incubated for >18 h to deplete residual carbon) or plastic monomers to generate second pre-cultures (starting OD_600_ = 0.1). Then, the second pre-cultures were inoculated into the main cultures at an initial OD_600_ of 0.1, 0.6 (for continuous fermentation) or 2.0 (for R-3HB production). *E. coli* strains were cultivated at 37 ℃, whereas *P. putida* strains were cultivated at 30 ℃.

## METHOD DETAILS

### Construction of plasmids and introduction of genetic modifications into *P. putida*

All plasmids constructed in this study were based on the suicide vector pEMG-RIS for genomic modification, and plasmid construction as well as genetic modification procedures followed the method described by Meng *et al*. [48]. All the oligonucleotides (Table S3) were ordered, and the sequencing work was done at Eurofins Genomics (Ebersberg, Germany).

### Online measurement of gas transfer rates

For online monitoring of the oxygen transfer rate (OTR), carbon dioxide transfer rate (CTR), and respiratory quotient (RQ) in 500 mL Erlenmeyer flasks, cultivations were performed in a Kuhner TOM shaker (Adolf Kühner AG, Birsfelden, Switzerland) with a filling volume of 25 mL. For the OTR measurements in 96-deepwell plates (2.5 mL), cultivations were carried out in a Kuhner microTOM shaker (Adolf Kühner AG) with filling volumes of 0.4 or 1 mL. All cultivations were conducted at 30 ℃ and 300 rpm with a 50 mm orbital throw.

### Growth Profiler Cultivation

For evaluating the genetic stability of the *P. putida* ETAB strain, cultures were cultivated in a Growth Profiler (Enzyscreen BV, Heemstede, Netherlands). Strains were grown in 24-half-deepwell microtiter plates (CR1424e; Enzyscreen) with a 2 mL working volume at 30℃ and 225 rpm using a 25 mm orbital throw. Biomass was monitored as the instrument-derived green value.

### Setup of continuous fermentation

The continuous fermentation was performed in a 1.3 L BioFlo 120 stirred-tank bioreactor (Eppendorf AG, Hamburg, Germany) with a working volume of 0.5 L, operated using DASware control software version 6.6. The bioreactor was equipped with online monitoring sensors, including a InPro 6800 Series O_2_ Sensor 12 mm (Mettler-Toledo AG, Switzerland) and a pH probe (EasyFerm Plus PHI K8 160) (Hamilton, Bonaduz, Switzerland), as well as a Pt100 temperature sensor. The pH was maintained at 7.0 ± 0.05 via automated addition of 1 M KOH. Off-gas was routed through a condenser and continuously analyzed for O_2_ and CO_2_ concentrations using a BlueVary gas sensor controlled by BlueVis software (v4.65; BlueSens Gas Sensor GmbH, Herten, Germany). Agitation was provided by a single six-blade Rushton turbine (53 mm diameter). During the batch phase, DO was controlled at 30% by varying the agitation speed between 200 and 1,200 rpm at a constant aeration rate of 0.5 vvm, with a maximum agitation speed of 600 rpm subsequently fixed for the continuous phase. Fresh medium was supplied during the continuous phase using an external Watson-Marlow 120U/DV peristaltic pump (Watson-Marlow, Spirax Group plc, UK) with a PharMed^®^ BPT biocompatible peristaltic pump tubing (0.8 mm I.D., 4.0 mm O.D., 1.6 mm wall thickness; Saint-Gobain Life Sciences – Bioprocess Solutions, Akron, OH, USA) at 50 mL h^−1^, corresponding to a dilution rate of 0.1 h^−1^, while culture volume was maintained via an overflow outlet positioned at the liquid surface. Foaming was suppressed by supplementation of the feed medium with 0.02% (v/v) Antifoam 204. Steady-state was considered achieved when CDW varied by less than 5% over five residence times (equivalent to the exchange of five working volumes).

### Transcriptome analysis and genome re-sequencing

For transcriptome analysis between the ETAB strain cells in steady state in medium 3 and medium 1 phase during continuous fermentation, 0.2 mL of culture was rapidly centrifuged (3,400× g, 1 min, 4 ℃), the supernatant was discarded, and the pellet was immediately resuspended in 0.5 mL RNAlater to stabilize RNA and stored at 4 ℃ overnight. For genome re-sequencing, selected isolates were cultivated overnight in LB medium, and 0.8 mL of culture was collected for genomic DNA extraction using the Monarch® Spin gDNA Extraction Kit (New England BioLabs). DNA concentrations were normalized to 50 ng μL^−1^ based on NanoDrop measurements (Thermo Scientific). All samples for transcriptome analysis and genome re-sequencing were processed by GENEWIZ (Azenta Life Sciences, Leipzig, Germany) for next-generation sequencing and bioinformatic analysis. The details of up- and downregulated genes are shown in Tables S4 and S5.

### Enzymatic plastic depolymerization and preparation of the cultivation medium

Enzymatic depolymerization of a blended polymer mixture (PET, PBAT, and TPU at equal mass) was performed in a 50 mL reactor with pH control as described in Branson *et al.* (2025) [32]. PET foil was purchased from Goodfellow GmbH. PBAT (BASF) has a monomer molar ratio of TA:AA:BDO = 0.222:0.278:0.500, whereas TPU (Soprema) has a molar ratio of fatty acid polyol:BDO:MDI = 1.4:1.0:2.1. Briefly, 1 g of polymer blend was suspended in a 20 mL reaction volume (5% w/v substrate load) and incubated for 72 h at 60 ℃ with PES-H1 FY (10 U g^−1^). The temperature was then reduced to 30 ℃ and the reaction was continued for 24 h after the addition of UMG-SP-2 (10 U g^−1^). The reaction was terminated by precipitation of the enzymes with acetonitrile.

For the preparation of the R-3HB production medium, acetonitrile was removed by overnight air flushing, and the resulting hydrolysate was diluted threefold and used as the carbon source to prepare the production medium as described above, followed by filtration through a 0.22 µm filter.

## QUANTIFICATION AND STATISTICAL ANALYSIS

### Quantification by HPLC-UV/RI

All analyses were conducted using an UltiMate 3000 HPLC system (Thermo Fisher Scientific, Waltham, MA, USA). For the quantification of L-lactic acid, (*R*)-3-hydroxybutyrate, adipic acid, ethylene glycol, 1,4-butanediol, glycolaldehyde, glycolate, glyoxylate and succinate, culture samples were centrifuged at 16,000× g for 5 min, and the supernatants were filtered through a 0.22 µm syringe filter prior to analysis. Separation was performed on an ion-exchange Metab-AAC column (300 × 7.8 mm, 10 µm; ISERA GmbH, Düren, Germany) using 5 mM H_2_SO_4_ as the mobile phase at a flow rate of 0.8 mL/min and a column temperature of 60 ℃. Analytes were detected using a UV detector at 210 nm in combination with a Shodex RI-101 refractive index detector maintained at 35 ℃ (Showa Denko, Tokyo, Japan).

Terephthalic acid and protocatechuate were quantified using a reversed-phase ISASpher 100-5 C18 BDS column (250 × 4.0 mm; ISERA) coupled to a UV detector (MWD-3000, Thermo Fisher Scientific) operating at 290 nm. The mobile phases consisted of acetonitrile (A) and 0.1% formic acid in water (B), with the following gradient program: 10% A/90% B for 2 min; linear increase to 99% A over 12 min and held for 2 min; decrease to 10% A over 2 min and maintained for 4 min. The column temperature was set to 40 ℃. Samples were prepared by mixing culture broth with acetonitrile at a 1:1 (v/v) ratio, and incubating at 4 ℃ for at least 4 h followed by centrifugation and filtration.

### Data analyses, model construction, simulation

Data analysis was performed using OriginPro 2021 (OriginLab Corporation, Northampton, MA, USA). All reported values are indicated as averages ± standard deviation. Significance analysis was performed by the determination of the standard deviation or standard error of the mean when indicated, followed by an ordinary one-way ANOVA with assumed Gaussian distribution, minimum p <0.05.

Simulations were performed using a curated version of the *P. putida* KT2440 iJN1463 genome-scale metabolic model [72]. The model was expanded to enable the utilization of polyester-derived monomers by completing the metabolic pathways for assimilation of terephthalate, ethylene glycol, adipic acid, and 1,4-butanediol. Missing reactions were added to ensure functional degradation routes toward central carbon metabolism, and all newly implemented reactions were mass- and charge-balanced. To represent the metabolic capabilities, specific non-native or thermodynamically infeasible routes were constrained during simulations. All modifications made to the model are available in the Table S6. Flux balance analysis was conducted using a sink reaction for R-3HB as the objective function. Polymer hydrolysate compositions were simulated by constraining the uptake rates of individual monomers according to experimentally defined polymer ratios. All code and models created in this work are available in the GitHub repository (github.com/AzizBenA/iJN1463_plastic_producer).

## RESOURCE AVAILABILITY

### Lead contact

Requests for further information and resources should be directed to and will be fulfilled by the lead contact, Lars M. Blank.

### Materials availability

Detailed information of all recombinant plasmids and genetically engineered strains in this study can be found in the supplementary material and will be shared by the lead contact upon request.

### Data and code availability

All data reported in this paper will be shared by the lead contact upon request.

### Author contributions

H.M., R.W., and L.M.B. conceived the project. H.M. designed and performed the experiments, and analyzed the data. T.K. designed the continuous fermentation, performed the experiment, and analyzed the data. A.B.A. re-constructed the metabolic model and assimilated the theoretical yield. A.S. and P.V. performed the experiment. H.M. wrote the manuscript. T.K., A.B.A., A.S., Y.B., T.S., H.B., U.T.B., R.W., and L.M.B. revised the manuscript. L.M.B. supervised the project and provided the funding for this work. All authors have approved the final version of the manuscript.

## Acknowledgments

We are especially grateful to Nick Wierckx for sharing the starting AB strain of this study. H.M. acknowledges his personal stipend from the China Scholarship Council (CSC). The study was financially supported by the European Union’s Horizon 2020 research and innovation program under grant agreement no. 870294 (MIX-UP project), and the Werner Siemens Foundation within the WSS project of the century catalaix. The laboratory of L.M.B. is partially funded by the German Research Foundation (DFG) under Germany’s Excellence Strategy within the Cluster of Excellence FSC 2186 “The Fuel Science Center” ID: 390919832.

## Declaration of interests

The authors declare no conflict of interest

## Supplemental information

Supplemental information associated with this article can be found online

## Notes

### Competing Interest Statement

The authors have declared no competing interest.

## References

1. Europe, P. (2025) Annual production of plastics worldwide from 1950 to 2024 Plastics Europe

2. OECD (2022) Global plastics outlook: policy scenarios to 2060, OECD Publishing, Paris

3. Hahladakis, J.N. (2020) Delineating and preventing plastic waste leakage in the marine and terrestrial environment. Environ. Sci. Pollut. Res. 27, 12830–12837

4. Yadav, V. et al. (2020) Framework for quantifying environmental losses of plastics from landfills. Resour. Conserv. Recycl. 161, 104914

5. Pottinger, A.S. et al. (2024) Pathways to reduce global plastic waste mismanagement and greenhouse gas emissions by 2050. Science 386, 1168–1173

6. Bachmann, M. et al. (2023) Towards circular plastics within planetary boundaries. Nat. Sustain. 6, 599–610

7. Wei, R. et al. (2020) Possibilities and limitations of biotechnological plastic degradation and recycling. Nat. Catal. 3, 867–871

8. Klauer, R.R. et al. (2024) Biological upcycling of plastics waste. Annu. Rev. Chem. Biomol. Eng. 15, 315–342

9. Xu, A. et al. (2023) Future focuses of enzymatic plastic degradation. Trends Microbiol. 31, 668–671

10. Wei, R. et al. (2025) Process insights for harnessing biotechnology for plastic depolymerization. Nat. Chem. Eng., 110–117

11. Tournier, V. et al. (2023) Enzymes’ power for plastics degradation. Chem. Rev. 123, 5612–5701

12. Ellis, L.D. et al. (2021) Chemical and biological catalysis for plastics recycling and upcycling. Nat. Catal. 4, 539–556

13. Tiso, T. et al. (2021) Towards bio-upcycling of polyethylene terephthalate. Metab. Eng. 66, 167–178

14. Liu, J. et al. (2021) Biodegradation and up-cycling of polyurethanes: Progress, challenges, and prospects. Biotechnol. Adv. 48, 107730

15. Stepnov, A.A. et al. (2024) Revisiting the activity of two poly(vinyl chloride)-and polyethylene-degrading enzymes. Nat. Commun. 15, 8501

16. Jendrossek, D. (2024) Polyethylene and related hydrocarbon polymers (“plastics”) are not biodegradable. N. Biotechnol. 83, 231–238

17. Liu, L. et al. (2025) Challenges of polyethylene (PE) biodegradation–a perspective. Biotechnol. Adv., 108717

18. Oiffer, T., et al. (2024) Chemo-enzymatic depolymerization of functionalized low-molecular-weight polyethylene. Angew. Chem. Int. Ed. 63, e202415012

19. Wei, R. and Bornscheuer, U.T. (2025) New biocatalytic approaches for plastic depolymerization. Engineering DOI: 10.1016/j.eng.2025.11.017

20. von Haugwitz, G., et al. (2023) Synthesis of modified poly(vinyl alcohol)s and their degradation using an enzymatic cascade. Angew. Chem. Int. Ed. 62, e202216962

21. Wei, R., et al. (2025) Standardization guidelines and future trends for PET hydrolase research. Nat. Commun. 16, 4684

22. Wei, R. et al. (2022) Mechanism-based design of efficient PET hydrolases. ACS Catal. 12, 3382–3396

23. Mican, J. et al. (2024) Exploring new galaxies: Perspectives on the discovery of novel PET-degrading enzymes. Appl. Catal. B-Environ. 342, 123404

24. Tournier, V. et al. (2020) An engineered PET depolymerase to break down and recycle plastic bottles. Nature 580, 216–219

25. Yoshida, S. et al. (2016) A bacterium that degrades and assimilates poly(ethylene terephthalate). Science 351, 1196–1199

26. Arnal, G. et al. (2023) Assessment of four engineered PET degrading enzymes considering large-scale industrial applications. ACS Catal. 13, 13156–13166

27. DeFrancesco, L. (2020) Closing the recycling circle. Nat. Biotechnol. 38, 665–669

28. Bell, E.L. et al. (2024) Natural diversity screening, assay development, and characterization of nylon-6 enzymatic depolymerization. Nat. Commun. 15, 1217

29. Bayer, T., et al. (2024) Structural elucidation of a metagenomic urethanase and its engineering towards enhanced hydrolysis profiles. Angew. Chem. Int. Ed. 63, e202404492

30. Branson, Y. et al. (2023) Urethanasen für die enzymatische hydrolyse niedermolekularer carbamate und das recycling von polyurethanen. Angew. Chem. 135, e202216220

31. Li, Z. et al. (2025) Discovery and engineering of a urethanase for enhanced depolymerization of polyurethane. ACS Catal. 15, 10768–10779

32. Branson, Y. et al. (2025) One-pot depolymerization of mixed plastics using a dual enzyme system. ChemSusChem 18, e202402416

33. Tiso, T. et al. (2022) The metabolic potential of plastics as biotechnological carbon sources–review and targets for the future. Metab. Eng. 71, 77–98

34. Valenzuela-Ortega, M. et al. (2023) Microbial upcycling of waste PET to adipic acid. ACS Cent. Sci. 9, 2057–2063

35. Sadler, J.C. and Wallace, S. (2021) Microbial synthesis of vanillin from waste poly(ethylene terephthalate). Green Chem. 23, 4665–4672

36. Narancic, T. et al. (2021) Genome analysis of the metabolically versatile *Pseudomonas umsongensis* GO16: the genetic basis for PET monomer upcycling into polyhydroxyalkanoates. Microb. Biotechnol. 14, 2463–2480

37. Parke, D. et al. (2001) Cloning and genetic characterization of *dca* genes required for β-oxidation of straight-chain dicarboxylic acids in *Acinetobacter* sp. strain ADP1. Appl. Environ. Microbiol. 67, 4817–4827

38. Ren, M. et al. (2025) NAD-dependent dehydrogenases enable efficient growth of *Paracoccus denitrificans* on the PET monomer ethylene glycol. Nat. Commun. 16, 5845

39. Op de Hipt, L., et al. (2025) Engineering of 1,4-butanediol and adipic acid metabolism in *Pseudomonas taiwanensis* for upcycling to aromatic compounds. Microb. Biotechnol. 18, e70205

40. Op de Hipt, L., et al. (2026) Process integration of enzymatic and microbial PBAT conversion with a *Pseudomonas taiwanensis* mixed culture. Syst. Microbiol. Biomanufacturing. 6, 16

41. Lee, H. et al. (2025) Developing an alternative medium for in-space biomanufacturing. Nat. Commun. 16, 728

42. Feng, C.-Q., et al. (2025) Screening and engineering of lycopene-producing strain *Rhodococcus jostii* for bio-upcycling of poly(ethylene terephthalate) waste. ScTEn 958, 178168

43. de Lorenzo, V., et al. (2024) *Pseudomonas putida* KT2440: the long journey of a soil-dweller to become a synthetic biology chassis. J. Bacteriol. 206, 1–14

44. Nikel, P.I. and de Lorenzo, V. (2018) *Pseudomonas putida* as a functional chassis for industrial biocatalysis: from native biochemistry to trans-metabolism. Metab. Eng. 50, 142–155

45. de Witt, J., et al. (2025) Upcycling of polyamides through chemical hydrolysis and engineered *Pseudomonas putida*. Nat. Microbiol., 1–14

46. Zobel, S. et al. (2015) Tn7-based device for calibrated heterologous gene expression in *Pseudomonas putida*. ACS Synth. Biol. 4, 1341–1351

47. Martínez-García, E. and de Lorenzo, V. (2011) Engineering multiple genomic deletions in Gram-negative bacteria: analysis of the multi-resistant antibiotic profile of *Pseudomonas putida* KT2440. Environ Microbiol 13, 2702–2716

48. Meng, H. et al. (2024) Establishing a straightforward I-SceI-mediated recombination one-plasmid system for efficient genome editing in *P. putida* KT2440. Microb. Biotechnol. 17, e14531

49. Volke, D.C. et al. (2022) Modular (de)construction of complex bacterial phenotypes by CRISPR/nCas9-assisted, multiplex cytidine base-editing. Nat. Commun. 13, 3026

50. Li, W.J. et al. (2019) Laboratory evolution reveals the metabolic and regulatory basis of ethylene glycol metabolism by *Pseudomonas putida* KT2440. Environ Microbiol 21, 3669–3682

51. Li, W.-J. et al. (2020) Unraveling 1,4-butanediol metabolism in *Pseudomonas putida* KT2440. Front. Microbiol. 11, 382

52. Ackermann, Y.S. et al. (2021) Engineering adipic acid metabolism in *Pseudomonas putida*. Metab. Eng. 67, 29–40

53. Werner, A.Z. et al. (2021) Tandem chemical deconstruction and biological upcycling of poly(ethylene terephthalate) to β-ketoadipic acid by *Pseudomonas putida* KT2440. Metab. Eng. 67, 250–261

54. Welsing, G. et al. (2025) Two-step biocatalytic conversion of post-consumer polyethylene terephthalate into value-added products facilitated by genetic and bioprocess engineering. Bioresour. Technol. 417, 131837

55. Diao, J. et al. (2025) Engineering microbial consortia for mixed plastic upcycling. Nat. Commun.

56. Bao, T. et al. (2023) Engineering microbial division of labor for plastic upcycling. Nat. Commun. 14, 5712

57. Utomo, R.N.C. et al. (2020) Defined microbial mixed culture for utilization of polyurethane monomers. ACS Sustainable Chem. Eng. 8, 17466–17474

58. Franden, M.A. et al. (2018) Engineering *Pseudomonas putida* KT2440 for efficient ethylene glycol utilization. Metab. Eng. 48, 197–207

59. Mückschel, B. et al. (2012) Ethylene glycol metabolism by *Pseudomonas putida*. Appl. Environ. Microbiol. 78, 8531–8539

60. Werner, A.Z. et al. (2025) Adaptive laboratory evolution and genetic engineering improved terephthalate utilization in *Pseudomonas putida* KT2440. Metab. Eng. 88, 196–205

61. Hara, H. et al. (2007) Transcriptomic analysis reveals a bifurcated terephthalate degradation pathway in *Rhodococcus* sp. strain RHA1. J. Bacteriol. 189, 1641–1647

62. Sasoh, M. et al. (2006) Characterization of the terephthalate degradation genes of *Comamonas* sp. strain E6. Appl. Environ. Microbiol. 72, 1825–1832

63. Köbbing, S. et al. (2024) Reliable genomic integration sites in *Pseudomonas putida* identified by two-dimensional transcriptome analysis. ACS Synth. Biol. 13, 2060–2072

64. Balusamy, K. et al. (2025) Continuous cultivation of defined mixed bacterial cultures for upcycling polyurethane and polyethylene terephthalate monomers to polyhydroxyalkanoates. Polym. Degrad. Stab., 111605

65. Espinosa-Urgel, M. and Ramos-González, M.I. (2023) Becoming settlers: Elements and mechanisms for surface colonization by *Pseudomonas putida*. Environ Microbiol 25, 1575–1593

66. Dmitrieva-Posocco, O. et al. (2022) β-Hydroxybutyrate suppresses colorectal cancer. Nature 605, 160–165

67. Kilpatrick, E.S. et al. (2023) Establishing pragmatic analytical performance specifications for blood beta-hydroxybutyrate testing. Clin. Chem. 69, 519–524

68. Yang, F.-R. et al. (2025) Hyperproduction of 3-hydroxybutyrate using engineered probiotic *E. coli* Nissle 1917 from glucose and CO_2_-derived acetate. Green Chem.

69. Manso Cobos, I., et al. (2015) *Pseudomonas pseudoalcaligenes* CECT5344, a cyanide-degrading bacterium with by-product (polyhydroxyalkanoates) formation capacity. Microb. Cell. Fact. 14, 77

70. Köbbing, S. et al. (2020) Characterization of context-dependent effects on synthetic promoters. Front. Bioeng. Biotechnol. 8, 551

71. Qian, Y., et al. (2026) A programmable microbial assembly line for plastic upcycling. Nat. Sustain., 1–13

72. Nogales, J. et al. (2020) High-quality genome-scale metabolic modelling of *Pseudomonas putida* highlights its broad metabolic capabilities. Environ Microbiol 22, 255–269

73. Eberz, J. et al. (2023) Selective separation of 4,4’-Methylenedianiline, Isophoronediamine and 2,4-Toluenediamine from enzymatic hydrolysis solutions of polyurethane. Solvent Extr. Ion Exch. 41, 358–373

74. Anderlei, T. et al. (2004) Online respiration activity measurement (OTR, CTR, RQ) in shake flasks. Biochem. Eng. J. 17, 187–194

75. Anderlei, T. and Büchs, J. (2001) Device for sterile online measurement of the oxygen transfer rate in shaking flasks. Biochem. Eng. J. 7, 157–162

76. Chacón, M. et al. (2024) Complex waste stream valorization through combined enzymatic hydrolysis and catabolic assimilation by *Pseudomonas putida*. Trends Biotechnol.

77. Lim, H.G. et al. (2022) Machine-learning from *Pseudomonas putida* KT2440 transcriptomes reveals its transcriptional regulatory network. Metab. Eng. 72, 297–310

78. Elmore, J.R. et al. (2020) Engineered *Pseudomonas putida* simultaneously catabolizes five major components of corn stover lignocellulose: Glucose, xylose, arabinose, p-coumaric acid, and acetic acid. Metab. Eng. 62, 62–71

79. Johnson, C.W. et al. (2017) Eliminating a global regulator of carbon catabolite repression enhances the conversion of aromatic lignin monomers to muconate in *Pseudomonas putida* KT2440. Metab. Eng. Commun. 5, 19–25

80. Wiegant, W.M. and De Bont, J.A.M. (1980) A new route for ethylene glycol metabolism in *Mycobacterium* E44. Microbiology 120, 325–331

81. Trifunović, D. et al. (2016) Ethylene glycol metabolism in the acetogen *Acetobacterium woodii*. J. Bacteriol. 198, 1058–1065

82. Lu, X. et al. (2019) Constructing a synthetic pathway for acetyl-coenzyme A from one-carbon through enzyme design. Nat. Commun. 10, 1378

83. Mainguet, S.E. et al. (2013) A reverse glyoxylate shunt to build a non-native route from C4 to C2 in *Escherichia coli*. Metab. Eng. 19, 116–127

84. Orsi, E. et al. (2025) Computation-aided designs enable developing auxotrophic metabolic sensors for wide-range glyoxylate and glycolate detection. Nat. Commun. 16, 2168

85. Monclús, L. et al. (2025) Mapping the chemical complexity of plastics. Nature 643, 349–355

86. Hartmans, S. et al. (1989) Metabolism of styrene oxide and 2-phenylethanol in the styrene-degrading *Xanthobacter* strain 124X. Appl. Environ. Microbiol. 55, 2850–2855

